# *Schistosoma japonicum* Histone Acetyltransferase 1 (*S*jHAT1): A Novel Anti-schistosomal Drug Target

**DOI:** 10.1101/2025.10.13.682244

**Authors:** Jing Xu, Yu-Xin Wang, Ping Huang, Ya-Nan Zhang, Huan Sun, Ting-Zheng Zhan, Chao-Ming Xia

**Affiliations:** Department of Parasitology, School of Basic Medical Sciences, Suzhou Medical College of Soochow University, No.199 Ren’ai Road, Suzhou City, Jiangsu Province, P. R. China; MOE Key Laboratory of Geriatric Diseases and Immunology, Suzhou Key Laboratory of Pathogen Bioscience and Anti-infective Medicine, Suzhou Medical College, Soochow University, Jiangsu Province, P. R. China; Key Laboratory of Basic Research on Regional Diseases (Guangxi Medical University), Education Department of Guangxi Zhuang Autonomous Region, Nanning City, Guangxi Province, P. R. China; Experimental Center of Suzhou Medical College, Soochow University, No.199 Ren’ai Road, Suzhou City, Jiangsu Province, P. R. China; The Second Affiliated Hospital of Soochow University, No.1055 Sanxiang Road, Suzhou City, Jiangsu Province, P. R. China

**Keywords:** *Schistosoma japonicum*, histone acetyltransferase 1, drug target

## Abstract

Schistosomiasis remains a critical global health issue, necessitating novel therapies due to emerging praziquantel resistance. We previously developed a patented praziquantel derivative, DW-3-15, which demonstrated potent broad-spectrum schistosomicidal activity through *Sj*HAT1 inhibition. Here, the full-length *Sj*HAT1 cDNA is cloned by rapid amplification of cDNA ends methods. The amino acids encoded by the cDNA retains conserved catalytic residues of acetyltransferases although sharing only 34% identity with mammalian orthologs. Phylogenetic analyses place it in a distinct clade, indicating its divergence specific to *Schistosoma* species. The expression profiling of *SjHAT1* reveals stage- and sex-specific patterns. Fluorescence in situ hybridization localizes *SjHAT1* predominantly in female vitellaria and male parenchyma near the gynecophoral canal. Knockdown of *SjHAT1* impairs worm survival, reduces female oviposition and disrupts ovarian and vitelline morphology both *in vitro* and *in vivo*. RNA-sequencing analysis reveals that *SjHAT1* knockdown disrupts β-alanyl-tryptamine pheromone signaling via downregulation of *aromatic L-amino acid decarboxylase* in males and *multidrug resistance-associated protein 4* in females. Through exploring this dual-sex regulatory mechanisms, *Sj*HAT1 may emerge as a promising therapeutic target for interrupting schistosomiasis transmission.

## 1. Introduction

Schistosomiasis is a neglected parasitic disease caused by infection with trematodes of the genus *Schistosoma*, mainly *Schistosoma mansoni*, *Schistosoma haematobium*, *Schistosoma japonicum*, *Schistosoma intercalatum*, *Schistosoma guineensis*, and *Schistosoma mekongi* (Buonfrate et al, 2025; World Health Organization, 2022). Current global data indicate that approximately 1 billion people are at risk of schistosomiasis, with more than 250 million people infected across 78 countries (Buonfrate et al, 2025; World Health Organization, 2022). Annually, about 250,000 deaths are attributed to this disease (Wang et al, 2020). In the absence of an effective vaccine, praziquantel (PZQ) is the only available drug to control this debilitating disease (McManus et al, 2020). However, PZQ is ineffective against immature worms and does not prevent reinfection, and repeated use of drug monotherapy in many endemic regions has raised serious concerns about the emergence of drug resistance (Berger et al, 2024; Summers et al, 2022). Therefore, developing alternative treatments and identifying novel drug targets for schistosomiasis remains crucial.

Currently, three strategies dominate the search for new anti-schistosomal drugs: (1) synthesis and evaluation of PZQ analogues, (2) design of new pharmacophores and (3) high-throughput screening of new active compounds (da Silva et al, 2017). While designing novel pharmacophores is challenging due to the still unclear mechanism of PZQ (Cupit & Cunningham, 2015), the development of PZQ derivatives remains a primary approach for discovering alternative therapeutics. In prior work, our group hybridized the endoperoxide bridge pharmacophore of artesunate (which targets juvenile schistosomes) to PZQ (which targets adults) by inserting a hydroxyl group at the position C10 of the pharmacophore aromatic ring in PZQ, yielding a novel praziquantel derivative DW-3-15 (Patent No. ZL201110142538.2) (Duan et al, 2012). DW-3-15 exhibited broad-spectrum schistosomicidal activity against all developmental stages of *S. japonicum in vivo*, with especially high efficacy against juveniles (Dong et al, 2014). The anti-juvenile and anti-adult effects of DW-3-15 were consistently observed in commercially synthesized batches, confirming its structural stability. Moreover, DW-3-15 significantly reduced egg deposition by female worms and mitigated hepatic pathology, further supporting its therapeutic potential (Wang et al, 2019). Notably, combining PZQ with DW-3-15 produced synergistic effects *in vitro* and *in vivo*; even at sub-therapeutic doses, the combination achieved efficacy comparable to or exceeding either monotherapy, suggesting the two compounds have distinct molecular targets (Yang et al, 2021).^]^

To investigate the molecular mechanism of DW-3-15, we performed tandem mass tag (TMT) quantitative proteomics on treated worms. This analysis revealed that *S. japonicum* histone acetyltransferase 1 (*Sj*HAT1) was one of the most significantly downregulated proteins in female worms following *in viv*o DW-3-15 treatment (Xu et al, 2024). HAT1 is a member of the GCN5-related N-acetyltransferases (GNAT) superfamily and is one of the first histone acetyltransferases discovered, primarily acetylating lysine 5 and lysine 12 of histone H4 (H4K5 and H4K12) (Fioravanti et al, 2020). HATs catalyze the transfer of acetyl groups to lysine residues on histones, thereby increasing chromatin accessibility to promote gene transcription (Park & Kim, 2020). This acetylation-mediated post-translational modification plays crucial roles in the growth, development, and reproduction of schistosomes (Carneiro et al, 2014; Cabezas-Cruz, 2014; Hong et al, 2016; Li et al, 2017). However, only few investigations on HAT inhibitors against schistosomes are currently available. Carneiro et al (2014) demonstrated that histone acetylation by *Sm*CBP1 and *Sm*GCN5 in *S. mansoni* establishes an epigenetic state necessary for *Smp14* gene activation and eggshell formation; the HAT inhibitor PU139 prevents chromatin decondensation at the *Smp14* promoter, thereby impairing egg production and disrupting female reproductive development (Carneiro et al, 2014). Furthermore, testing of the HAT inhibitors A485, C646, and curcumin identified curcumin as a potent schistosomicidal agent that dose-dependently suppresses *Sj*HAT1 expression. Notably, we also found that DW-3-15 could inhibit *Sj*HAT1 expression, implicating *Sj*HAT1 as a promising target mediating its anti-schistosomal effects (Xu et al, 2024).

Despite these findings, the biological roles of *Sj*HAT1 in *S. japonicum*, particularly its involvement in drug-mediated killing of worms and in regulating female reproductive activity, remain poorly characterized. In this study, we employed rapid amplification of cDNA ends (RACE) to systematically analyze the bioinformatics features of *Sj*HAT1. Subsequently, we investigated its functions both *in vitro* and *in vivo*. Finally, we utilized RNA sequencing (RNA-seq) to explore the regulatory role of *Sj*HAT1 in downstream signaling pathways, thereby establishing experimental and theoretical foundations for the development of *Sj*HAT1-targeted anti-schistosomal interventions.

## 2. Results

### 2.1. *Sj*HAT1 is a promising drug target against schistosomiasis

Since the cDNA sequence of *SjHAT1* in the NCBI database is predicted and incomplete, we utilized the rapid amplification of cDNA ends (RACE) technique to obtain its full-length cDNA sequence for further characterization. The full-length cDNA sequence of *SjHAT1* was successfully obtained through RACE assay, revealing a 1302 bp open reading frame (ORF) encoding 433 amino acids (NCBI accession no. PQ778043). Using the InterPro database, we identified that *Sj*HAT1 contains an N-terminal HAT1 domain (IPR019467, 14-175aa) and a C-terminal HAT type B catalytic subunit domain (IPR048776, 316-339aa). The N-terminal HAT1 domain catalyzes the transfer of an acetyl group from acetyl-CoA to lysine residues at the histone N-terminus (Neuwald & Landsman, 1997). By contrast, the C-terminal domain is not conserved and is dispensable for HAT1 catalytic activity (Wu et al, 2012). To determine the enzymatic activity of *Sj*HAT1, we expressed recombinant *Sj*HAT1 in *E. coli* and found that it exhibits significant histone acetyltransferase activity *in vitro* (Fig. EV1 and Table EV1). To validate whether *Sj*HAT1 has potential as a drug target, we performed a multiple sequence alignment of *Sj*HAT1 with HAT1 homologs from *Homo sapiens*, *Mus musculus*, *Rattus norvegicus*, *Schistosoma mansoni*, *Schistosoma haematobium*, *Clonorchis sinensis*, and *Fasciola hepatica*. Notably, multi-sequence alignment revealed that *Sj*HAT1 shares only ∼34% sequence identity with the HAT1 proteins of human, mouse, and rat, although the key amino acids required for HAT1 enzymatic activity are highly conserved (Fig. 1A). Furthermore, phylogenetic analysis confirmed that *Sj*HAT1 clusters in an evolutionarily distant clade (Fig. 1B). These distinctive molecular characteristics, particularly the structural divergence from mammalian orthologs, establish *Sj*HAT1 as a promising druggable target for schistosomiasis. Such divergence suggests that species-specific targeting of *Sj*HAT1 could mitigate host toxicity by avoiding interference with the host’s HAT1.

**Figure 1.**
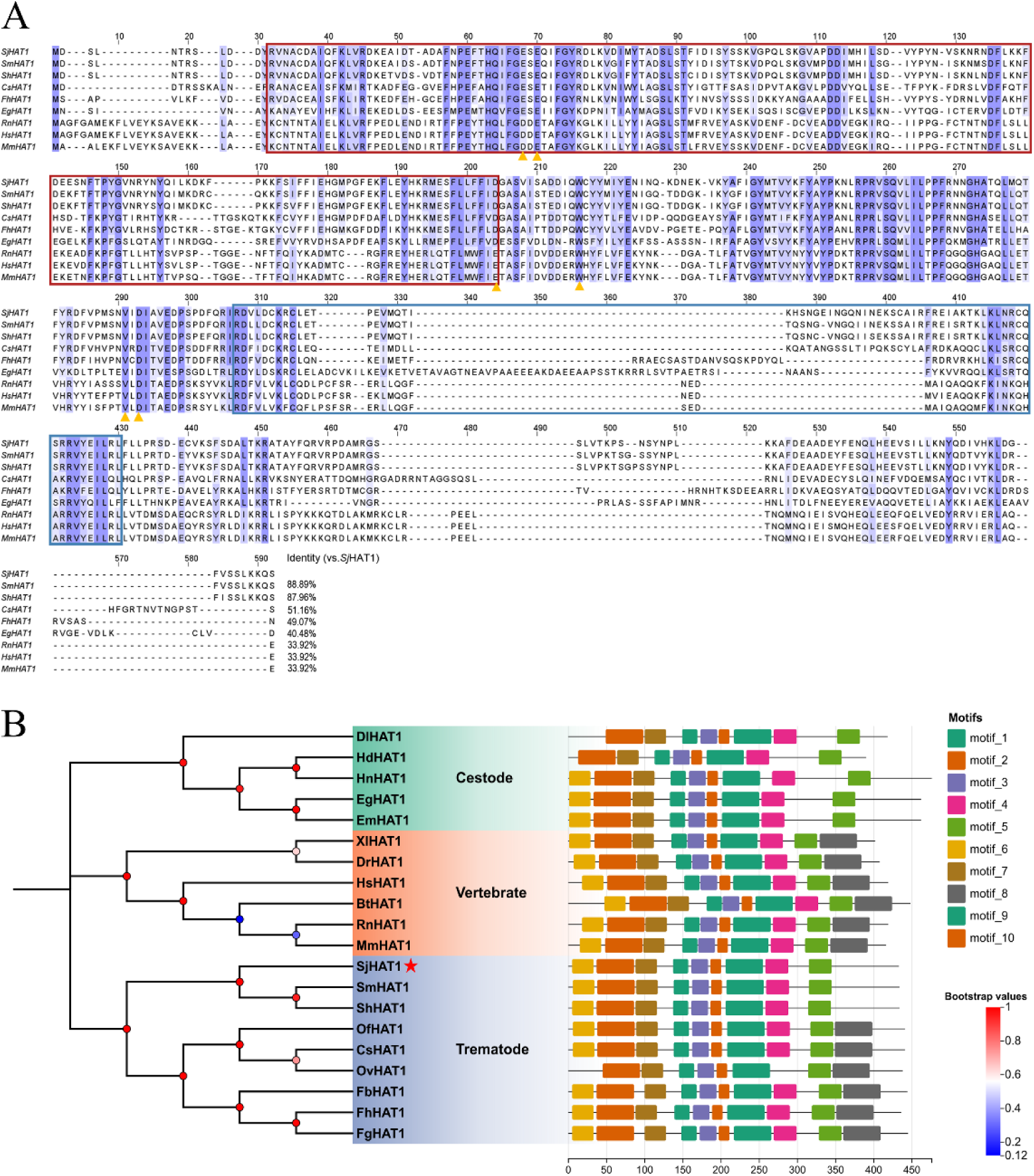
Multiple sequences alignment and phylogenetic analysis of *Sj*HAT1. (**A**) Multiple sequences alignment of *Sj*HAT1. Red box indicates the N-terminal domain of HAT1, blue box indicates the C-terminal domain of HAT1, and yellow triangles highlight key amino acids experimentally validated to be associated with HAT1 protein activity in other species (**B**) phylogenetic analysis of *Sj*HAT1. Red star indicates HAT1 in *S. japonicum*.

### 2.2. *SjHAT1* is highly expressed in the vitellaria of adult female worms

To further characterize *Sj*HAT1 and facilitate future investigations into its physiological functions, the mRNA expression profiles of the *SjHAT1* across six critical developmental stages, including eggs, cercariae, 14-day-old juveniles, 21-, 28- and 35-day-old adult male/female worms were quantified. The results revealed that *SjHAT1* transcripts, while ubiquitously expressed, exhibited significant stage specificity and sexual dimorphism. The highest expression level of *SjHAT1* was observed in 35-day-old females, whereas the lowest expression was detected in eggs. Notably, sex-specific differences became pronounced in 28- and 35-day-old adults, with *SjHAT1* expression in females significantly exceeding that in males (*p* < 0.0001, Fig. 2A). This differential expression pattern suggests that *SjHAT1* may play a pivotal role in sexual maturation and female reproductive development in schistosomes. Subsequently, we investigated the tissue-specific localization of *SjHAT1* in 35-day-old adult worms. Fluorescence *in situ*hybridization (FISH) revealed that *SjHAT1* is predominantly expressed in the vitellaria of female worms (Fig. 2B) and in the parenchymal cells adjacent to the gynecophoral canal of male worms (Fig. 2C). The female-biased expression pattern in vitellaria, a specialized organ responsible for vitellocytes production(Wang & Collins, 2016), further implicates *Sj*HAT1 in reproductive regulation.

**Figure 2.**
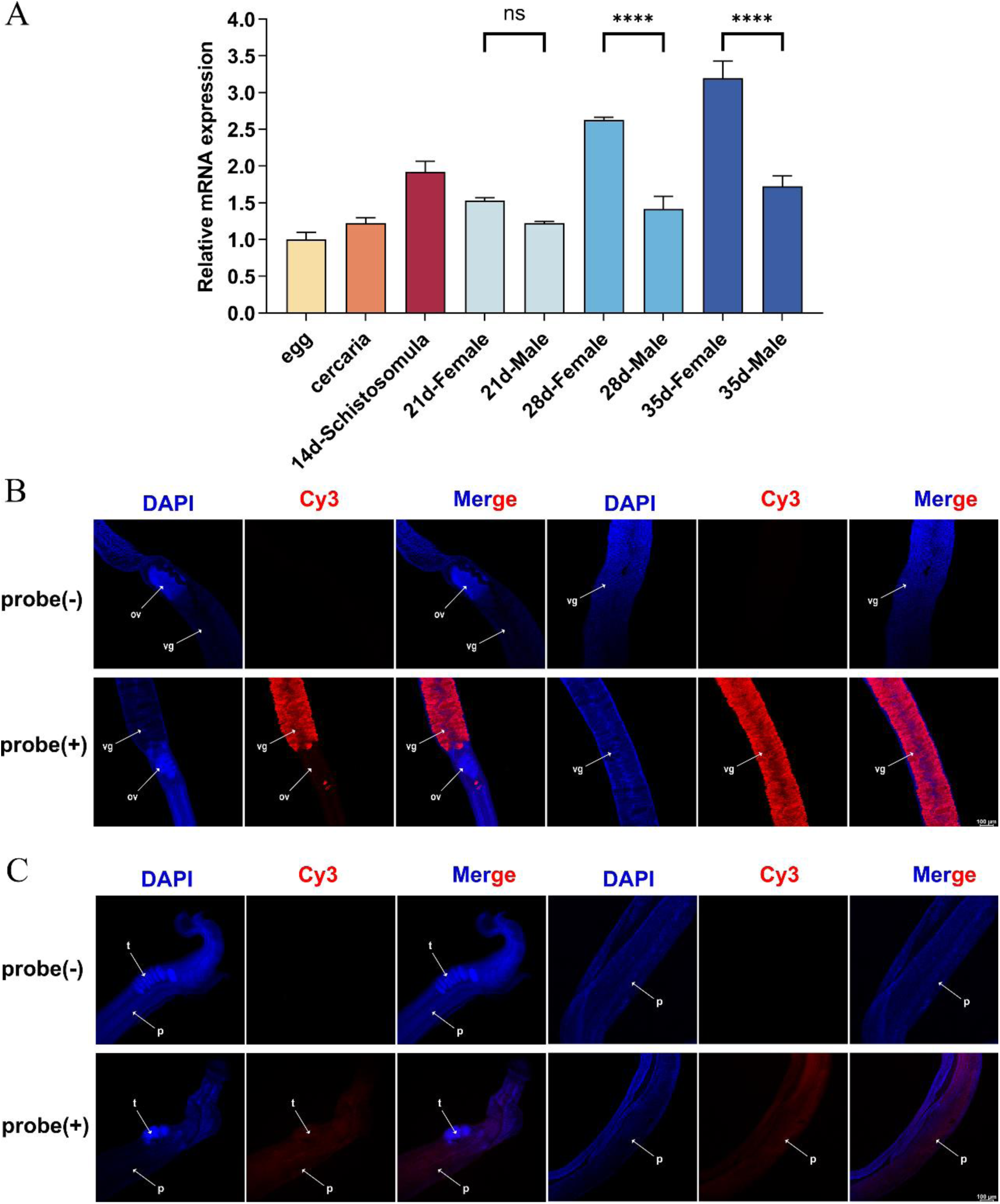
Expression and location of *Sj*HAT1 in adult worms of *S. japonicum*. (**A**) Transcriptional levels of *SjHAT1* in different developmental stages and genders of *S. japonicum*. N = 3 biological replicates, One-Way ANOVA, standard deviation (SD) is shown in the error bars. ‘ns’ indicates no significant difference (*p* > 0.05), \*\*\*\**p* < 0.0001. (**B**) Localization of *SjHAT1* in female *S. japonicum*. ov, ovary; vg, vitelline gland. Scale bars: 200 μm. (**C**) Localization of *SjHAT1* in male *S. japonicum*. t, testis; p, parenchymal cell. Scale bars: 200 μm.

### 2.3. *SjHAT1* knockdown impairs viability and oviposition of adult *S. japonicum in vitro*

To investigate the functions of *Sj*HAT1, we conducted RNA interference (RNAi) in paired adult worms *in vitro*. We designed two dsRNAs, *SjHAT1*-dsRNA1 and *SjHAT1*-dsRNA2, targeting the *N*-terminal and *C*-terminal domains of *Sj*HAT1, respectively. Through preliminary experiments comparing the interference efficiency of *SjHAT1*-dsRNA1 (30 μg/mL), *SjHAT1*-dsRNA2 (30 μg/mL), and *SjHAT1*-dsRNA1/2 (15 μg/mL *SjHAT1*-dsRNA1 and 15 μg/mL *SjHAT1*-dsRNA2), we found that the *SjHAT1*-dsRNA1/2 exhibited the highest interference efficiency (Fig. EV2). Hence, in the following studies, *SjHAT1*-dsRNA1/2 was added on day 1, day 3, and day 7 using the soaking method (Wang et al, 2020). The results showed that *in vitro* RNAi significantly reduced both the transcriptional level and protein expression of the *Sj*HAT1 in female and male worms (Fig. 3A-B).

**Figure 3.**
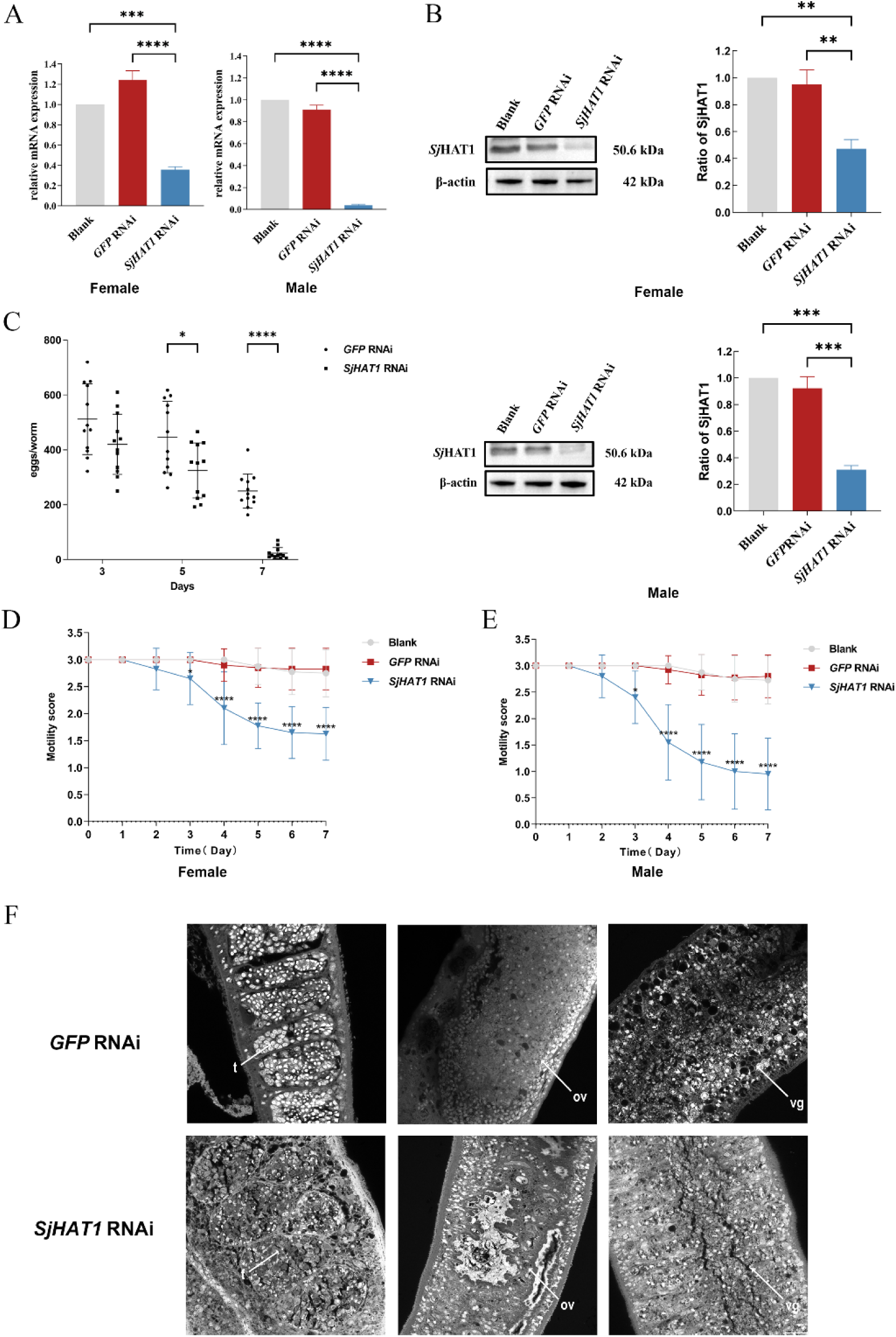
*SjHAT1* knockdown disrupts the viability and oviposition in adult worms *in vitro*. (**A**) Changes in *SjHAT1* transcript level after *SjHAT1*-dsRNA treatment. Standard deviation (SD) is shown in the error bars. N= 3 bilogical replicates, One-way ANNOVA, \*\*\*\**p* < 0.0001. (**B**) Changes in *Sj*HAT1 protein expression after *SjHAT1*-dsRNA treatment. Standard deviation (SD) is shown in the error bars. N= 3 bilogical replicates, One-way ANNOVA, \*\**p* < 0.01, \*\*\**p* < 0.001. (**C**) Altered oviposition in females after *SjHAT1*-dsRNA treatment. N=12, Two-Way ANOVA, \**p* < 0.05, \*\*\*\**p* < 0.0001. (**D**) Altered Motility in females after *SjHAT1*-dsRNA treatment. N=40, Two-Way ANOVA, \*\*\*\**p* < 0.0001. (**E**) Altered Motility in males after *SjHAT1*-dsRNA treatment. N=40, Two-Way ANOVA, \*\*\*\**p* < 0.0001. (**F**) Morphological changes in the reproductive organs of worms after *SjHAT1*-dsRNA treatment. t, testis; ov, ovary; vg, vitelline gland. Scale bars: 20 μm. Standard deviation (SD) is shown in the error bars.

After 7 days of *SjHAT1*-dsRNA treatment, *S. japonicum* adult worms exhibited pronounced phenotypic alterations, including markedly reduced egg laying by females and compromised locomotor activity. As shown in Fig. 3C, female oviposition was significantly reduced at day 5 posttreatment with *SjHAT1*-dsRNA interference (*p*<0.05), reaching the lowest at day 7 (*p*<0.0001). Similarly, both female and male worms exhibited significantly lower viability compared with the control group from day 3 onward (Fig. 3D-E, *p*<0.05), with scores reaching their lowest levels by 7 days posttreatment (Fig. 3D-E, *p*<0.0001). At day 7, the intervention showed no statistically significant impact on worm pairing (Fig. EV3).

To assess reproductive organ development following *SjHAT1* knockdown, worms were carmine-stained and examined by confocal laser scanning microscopy (CLSM). We found that knockdown of *SjHAT*1 severely disrupted the morphology of the reproductive organs in both sexes of *S. japonicum*. In males, the testes became disorganized, with indistinct lobular boundaries, loose stromal matrices, reduced spermatogonial populations, and abnormal cellular morphology. In females, the vitelline glands were atrophic, characterized by effaced lobular boundaries, fewer mature vitelline cells, and markedly shrunken ovaries (Fig. 3F).

It is well known that an intact tegument is crucial for worm survival (da Silva et al, 2017). Disruption of the tegument leads to exposure of the worm’s antigens, rendering it more susceptible to the host immune system (Vale et al, 2017; Xu et al, 2023). We therefore employed scanning electron microscopy (SEM) to examine the tegumental changes after *SjHAT*1 knockdown. According to the *in vitro* experimental results, worm viability began to decline significantly by 3 days post-interference. Consequently, parasites collected on the third day after interference were subjected to SEM observation. Under SEM, the tegument of the female worm in the control group remained intact, with spines uniformly arranged in the ventral sucker (Fig. EV4). In contrast, following *SjHAT1*-dsRNA intervention for 3 days, distinct tegumental damage was evident. The anterior region of female worms exhibited mild swelling accompanied by numerous blisters (Fig. 4B). The mid-body tegument displayed extensive sloughing (Fig. 4C), and obvious fusion of the spines occurred in the ventral suck of females (Fig. 4D). These *in vitro* results indicate that *SjHAT1* knockdown critically impairs survival and reproductive functions in adult *S. japonicum*.

**Figure 4.**
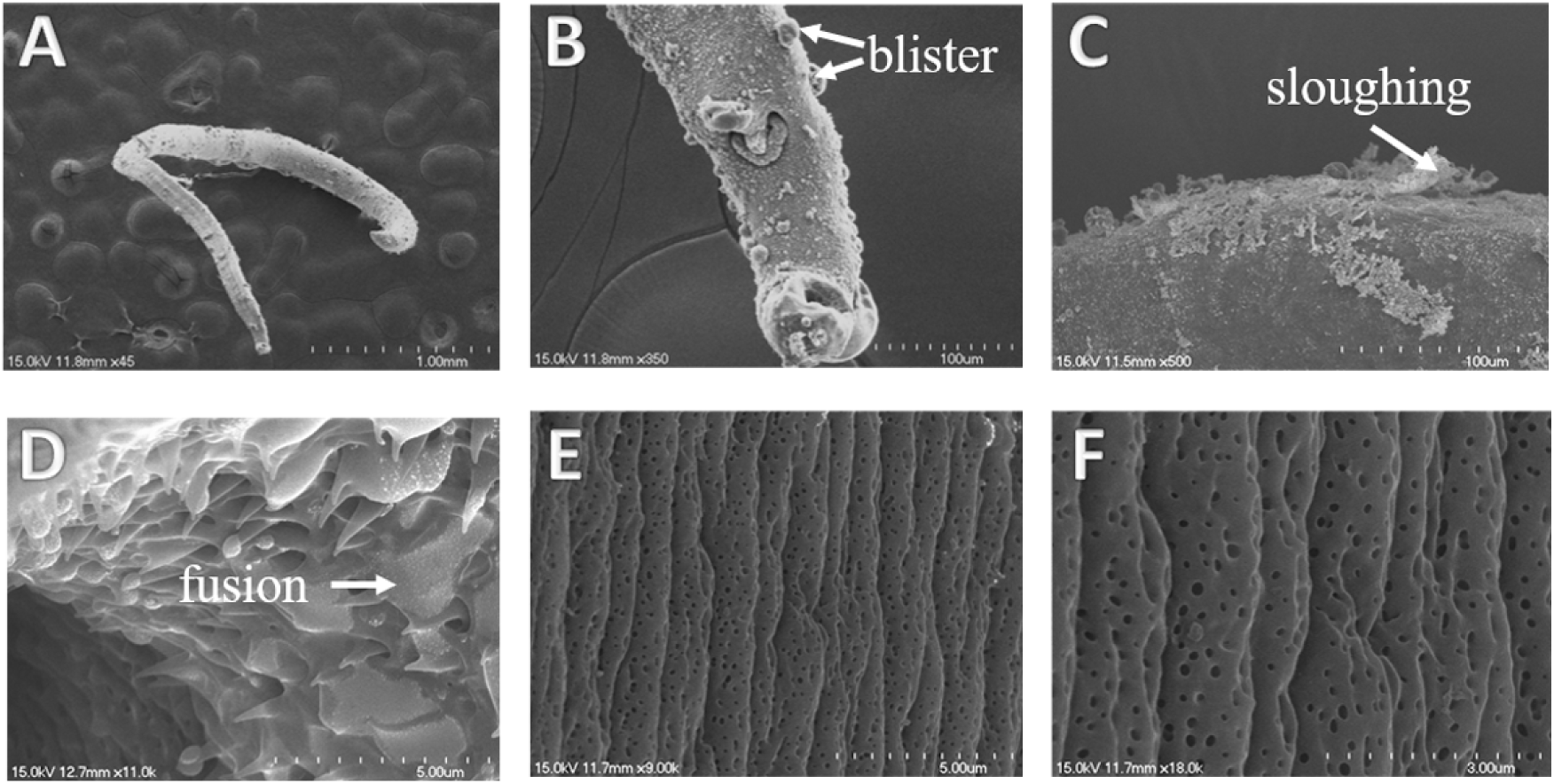
Scanning electron micrographs of the tegument of female *S. japonicum* worms after *SjHAT1* RNAi for 3 days. (A)The whole worm is swollen and shrunken; (B) mild swelling accompanied by numerous blisters is observed at the anterior end of female worm; (C)extensive sloughing is observed on the tegument in the mid-body of female worm; (D) fusion of the spines in the ventral sucker are obvious; (E)-(F) no significant changes are observed in the transverse ridge-like structures on the tegument of female worm. The ridges arranged orderly accompanied by uniformly distributed pores. Scalebars: A: 1 mm; B: 100 μm; C: 100 μm; D: 5 μm; E: 5 μm; F: 3 μm.

### 2.4. *SjHAT1* is essential for the survival, pairing and oviposition of *S. japonicum in vivo*

Our *in vitro* results indicate *Sj*HAT1 is required for maintaining the adult worm viability and is crucial for reproductive processes in *S. japonicum*. However, the *in vitro SjHAT1*-dsRNA interference experiment was conducted on paired adult worms and did not address the role of *Sj*HAT1 at other developmental stages. Furthermore, it is technically challenging to collect and culture schistosomula of different developmental stages *in vitro*. Thus, we carried out *in vivo* RNAi experiments to assess the effects of *SjHAT1* knockdown across parasite developmental stages. Mice infected with *S. japonicum* cercariae received a tail vein injection of 10 μg *SjHAT1*-dsRNA at 14 days post-infection (dpi), followed by repeated injections of 10 μg dsRNA at 18, 22, 26, and 30 dpi. On day 35 post-infection, the parasites were harvest through perfusion of the hepatic portal system and mesenteric veins, and mouse livers were collected for egg counting and histopathological analysis (H&E staining). Worm burden reduction and hepatic egg burden reduction were calculated.

In these experiments, mice that received control *GFP*-dsRNA showed no significant differences in worm or egg burdens compared to blank controls injected with saline (Fig. EV5 A-B and 5E). Parasites recovered from *GFP*-dsRNA control mice were morphologically normal (Fig. EV5C-D), and there were no significant differences in the number or size of liver egg granulomas between *GFP*-dsRNA and blank control groups (Fig. EV5F-G). These findings confirm that *GFP*-dsRNA had no appreciable effect on the parasites. In contrast, mice treated with *SjHAT1*-dsRNA showed a 57.5% reduction in worm burden (Fig. 5A, p<0.0001) and a 65.9% reduction in pairing (Fig. 5B, p<0.0001) compared to the *GFP*-dsRNA controls. Moreover, many female worms from the *SjHAT1* knockdown group exhibited developmental arrest (Fig. 5C), with 32.9% remaining immature, which was significantly higher than that the 18.9% observed in the *GFP* control group [*χ*^2^(1) =5.528, *p*=0.0187, Table EV2]. These results indicate that the downregulation of *SjHAT1* impairs the development and sexual maturity of female worms, thereby reducing their egg-laying capacity. Consistent with these observations, CLSM analysis revealed severe testicular hypoplasia in males, while females showed ovarian degeneration and vitelline gland atrophy characterized by diminished lobular boundaries and reduced mature vitellocyte count (Fig. 5D). Eggs are the key factor responsible for the pathogenesis of schistosomiasis (Popiel, 1986; Kunz, 2001; LoVerde,2002; Sun et al, 2023). The formation of hepatic egg granulomas depends not only on the number of deposited eggs but also on their viabilities. Pathologically, *SjHAT1*-RNAi livers displayed significantly attenuated hepatic granuloma formation (Fig. 5E) and 79.2% reduction in egg deposition (*p*<0.0001, Fig. 5F). H&E staining confirmed 82.9% smaller granuloma areas 9480±5029 μm^2^ vs control 55542±17398 μm^2^ (*p*<0.0001, Fig. 5G-H), demonstrating impaired oviposition capacity post-RNAi. These findings indicate that *SjHAT1* knockdown *in vivo* severely impairs oviposition and significantly mitigates egg-induced hepatic pathology.

**Figure 5.**
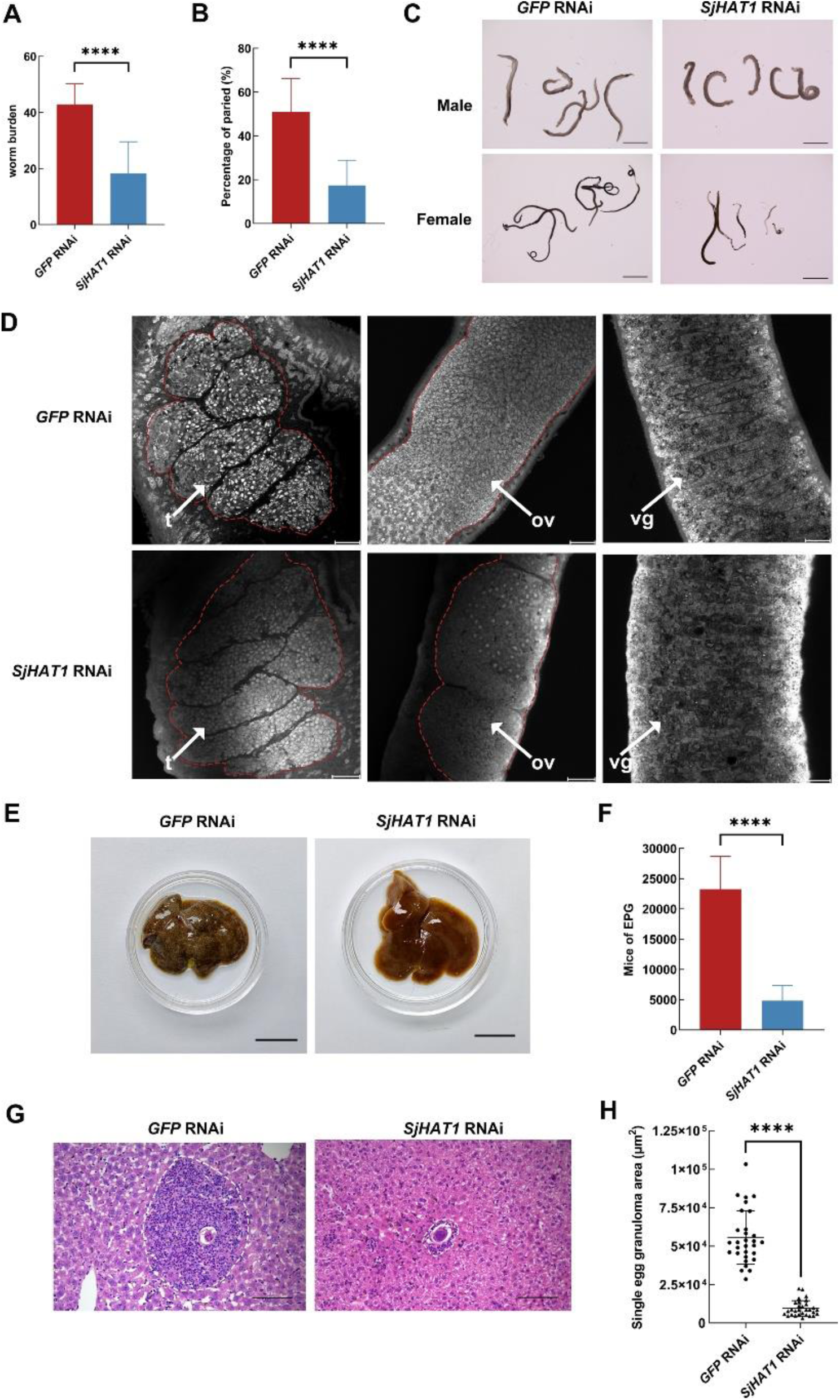
*Sj*HAT1 affects survival, pairing and oviposition of *S. japonicum in vivo*. On the 14^th^, 18^th^, 22^th^, 26^th^ and 30^th^ days postinfection, *SjHAT1*-dsRNA was injected through the tail vein, and the hepatic portal vein was perfused on the 35^th^ day. (A) Worm burden of the parasites recovered at 35 dpi in *GFP* control RNAi and *SjHAT1* RNAi groups. N = 3 biological replicates, 3 mice of each biological replicates. Student’s *t*-test, \*\*\*\**p* < 0.0001. (B) Comparation of the pairing rates of recovered worms between *GFP* and *SjHAT1* dsRNA groups. N = 3 biological replicates, 3 mice of each biological replicates. Student’s *t*-test, \*\*\*\**p* < 0.0001. (C) Morphological observation of worms after RNAi *in vivo*. Scale bars: 2000 μm. (D) Morphological changes in the reproductive organs of worms after RNAi *in vivo*. t, testis; ov, ovary; vg, vitelline gland. Scale bars: 20 μm. (E) Gross observations of the mouse liver in the *GFP* and *SjHAT1* dsRNA groups. Scale bars: 1 cm. (F) Eggs per gram of the liver comparation between the *GFP* and *SjHAT1* dsRNA groups (n=90). Student’s *t*-test, \*\*\*\**p* < 0.0001. (G) Histological assessment of mouse liver by H&E staining. Scale bars: 100 μm. (H) Statistical analysis of the size of egg granuloma area after RNAi *in vivo.* N=30, standard deviation (SD) is shown in the error bars. Student’s *t*-test, \*\*\*\**p* < 0.0001.

### 2.5. *SjHAT*1 knockdown modulates ABC transporter pathway in females and tyrosine metabolism in males

Having demonstrated that *SjHAT1* knockdown disrupts the normal morphology, growth, development and oviposition of *S. japonicum*, we next investigated the molecular pathways regulated by *SjHAT1*. We performed RNA sequencing (RNA-seq) analysis on male and female worms following *SjHAT1* knockdown *in vitro*. The RNA-seq results revealed a total of 206 differentially expressed genes (DEGs) in female worms (86 upregulated and 120 downregulated; Fig. 6A) and 786 DEGs in male worms (242 upregulated and 584 downregulated; Fig. 6B). Gene Ontology (GO) analysis and KEGG pathway enrichment were then applied to these DEGs.

**Fig. 6.**
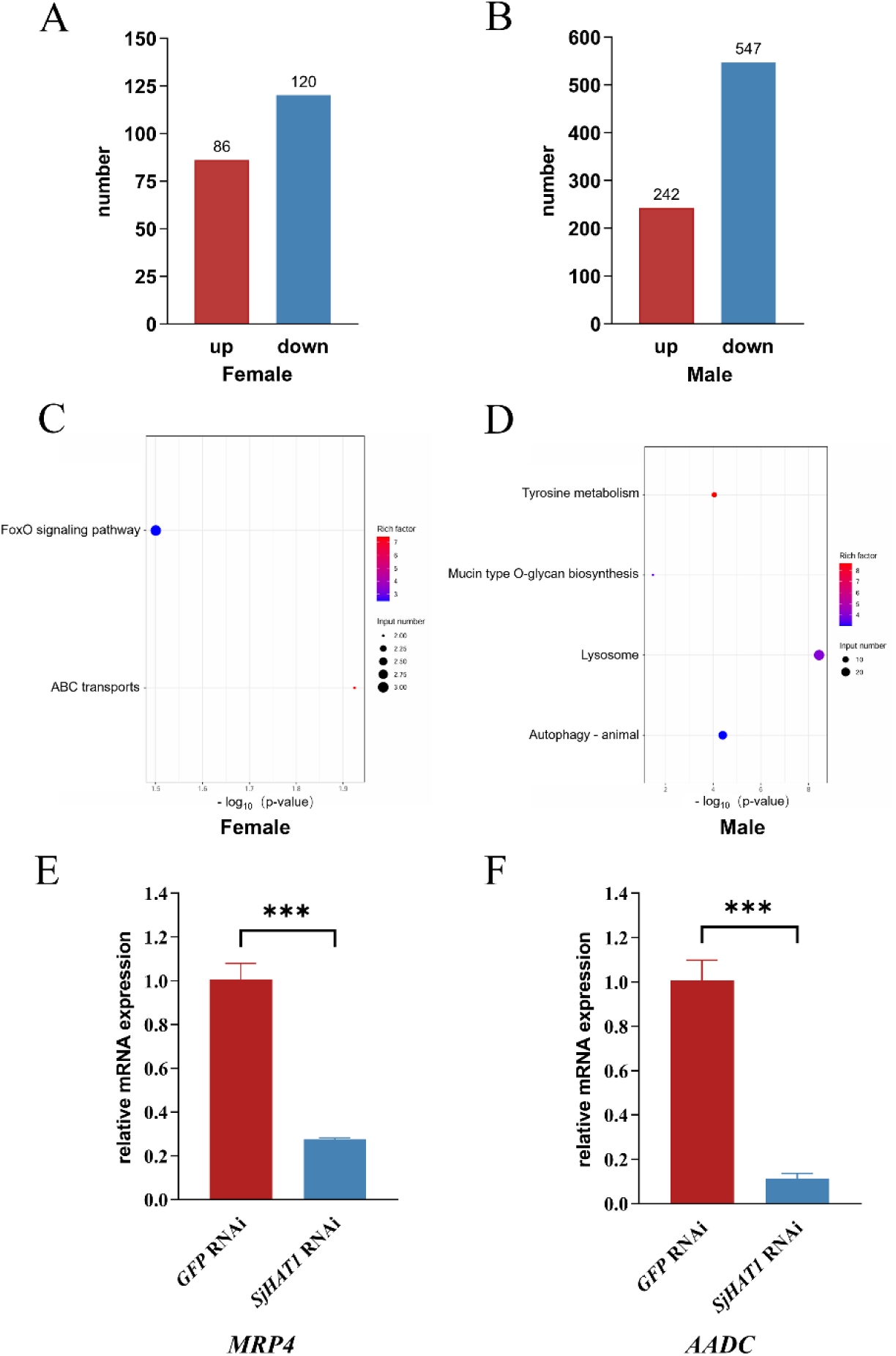
*Sj*HAT1 regulates different pathways in female and male worms. (A) Bar plot illustrating differentially expressed genes (DEGs) in female parasites. (B) Bar plot illustrating DEGs in male worms. DEGs were defined as |log2 Fold Change| ≥0.58 and *p*-value< 0.05. (C) KEGG pathway enrichment analysis of DEGs in female worms. (D) KEGG pathway enrichment analysis of DEGs in male worms. (E) qRT-PCR validation of *MRP4* expression in female worms. (F) qRT-PCR validation of *AADC* expression in male worms. Standard deviation (SD) is shown in the error bars. Differences are statistically significant (\*\*\**p* < 0.001, Student’s *t*-test).

After the knockdown of *SjHAT1*, genes with the most significant expression changes included *copine-9* (KSF78_0001940), *calcium activated potassium channel subunit* (KSF78_0002097), *tyrosine-protein kinase FRK* (KSF78_0003289), *ribonuclease T2* (KSF78_0002281), *multidrug resistance-associated protein 4* (*MRP4*, KSF78_0000799) and *multidrug resistance protein 1* (KSF78_0005913) in females, and *neuronal acetylcholine receptor subunit alpha-2* (KSF78_0006978), *aromatic L-amino acid decarboxylase* (*AADC*, KSF78_0006508), *tyrosinase* (KSF78_0005796, KSF78_0005182), *tyrosine hydroxylase* (KSF78_0002665) in males.

In female worms, the downregulated DEGs were significantly enriched in the ATP binding cassette (ABC) transporter pathway (Fig. 6C), with *MRP4* being specifically and significantly downregulated (*p* = 0.01931655). Consistently, GO analysis linked this female DEG to transmembrane transport processes (GO:0055085). In male worms, the downregulated DEGs were predominantly enriched in the tyrosine metabolism pathway (Fig. 6D). Notably, *AADC*, a gene normally highly expressed in males, was dramatically downregulated (*p* = 5.49458E-07). GO analysis indicated that this enzyme is involved in carboxylic acid metabolic processes (GO:0019752). Finally, qRT-PCR validation confirmed that *SjHAT1* knockdown significantly reduced *MRP4* expression in female worms (Fig. 6E) and *AADC* transcriptional levels in male worms (Fig. 6F), consistent with the RNA-seq data.

## 3. Discussion

In our previous work, we observed that the praziquantel derivative DW-3-15, has exceptional schistosomicidal activity against all developmental stages of *S. japonicum*, with >60% worm reduction against juveniles (Dong et al, 2014). This broad efficacy contrasts sharply with PZQ, which mainly targets adult worms (Xiao et al, 1987). Notably, DW-3-15 was found to markedly suppress the expression of *Sj*HAT1 in adult worms. Similarly, curcumin, a histone acetyltransferase inhibitor that also suppresses *Sj*HAT1 expression, elicited phenotypic changes resembling those caused by DW-3-15 (Xu et al, 2024). These findings suggest that *Sj*HAT1 is a critical molecular target underlying the anti-schistosomal effects of DW-3-15. However, the existing *SjHAT1* cDNA sequence in the NCBI database is incomplete and based on predictions. We used RACE to clone the full-length *SjHAT1* cDNA (NCBI accession no. PQ778043) and characterized the encoded protein. We identified four nucleotide differences from the predicted sequence, including a nonsynonymous mutation (cysteine to serine) at residue 311. Phylogenetic analysis confirmed that *Sj*HAT1 is highly diverged from its mammalian orthologs (*Hs*HAT1 in humans, *Mm*HAT1 in mice), reflecting their distant evolutionary relationship (Fig. 1B). This divergency implies that pharmacological agents could be designed to specifically target *Sj*HAT1’s unique domains, potentially minimizing off-target effects on homologous host proteins and thereby mitigating toxicity. Importantly, our qPCR analysis confirmed constitutive expression of *SjHAT1* across all developmental stages and both genders of *S. japonicum* (Fig. 2A), supporting its essential biological role throughout the parasite’s life cycle. Its expression level gradually increased as the parasite matured, with significantly higher expression observed in mature female worms compared to their male counterparts of the same age. This sex-biased expression pattern suggests that *Sj*HAT1 may play a critical role in the development and maturity of the female worms. Consistent with the expression data, FISH revealed that *SjHAT1* is predominantly localized in the vitellaria of female *S. japonicum* (Fig. 2B). The vitellaria occupy approximately two-thirds of the female body and comprise numerous transversely arranged vitelline lobules extending dorsoventrally. These lobules contain vitelline cells at various developmental stages, which provide both nutrients for fertilized ova and polyphenolic proteins critical for eggshell formation (MOORE & SANDGROUND, 1956; He et al, 1974; Wang et al, 2016; Sun et al, 2023). Mature female schistosomes produce an average of 3,500 eggs per day. Each mature egg consists of one fertilized ovum surrounded by approximately 20 vitelline cells, requiring the female to synthesize 70,000 vitelline cells daily (equivalent to 50 cells per minute) to sustain oviposition (He et al,1974). Disruption of vitelline gland function would directly impair egg production. Targeting egg-laying mechanisms not only mitigates immunopathological damage in the host but also represents a strategic intervention for controlling schistosomiasis transmission (Sun et al, 2023).

Our *in vitro* experiments further reinforce the potential of *Sj*HAT1 as a promising druggable target for schistosomiasis. Specifically, dsRNA-mediated silencing of *SjHAT1* significantly reduced schistosome viability and impaired egg-laying capacity in females, exhibiting time-dependent inhibitory effects. Morphological analysis indicated that *SjHAT1*-dsRNA impairs the integrity of the female tegument (Fig. 4C), a structure critical for schistosome survival (da Silva et al, 2017; Vale et al, 2017; Xu et al, 2023). Moreover, *SjHAT1* knockdown induced notable structural changes in the reproductive organs of *S. japonicum* adult worms. Following RNA interference, female ovaries displayed severe atrophy and distinct morphological abnormalities, including a marked reduction in mature oocytes, disappearance of intercellular spaces, disorganized cell arrangement, and vacuole formation (Fig. 3F). These phenotypic alterations were consistent with those induced by the HAT inhibitor PU139 (Carneiro et al, 2014). A previous study also demonstrated that HAT1 was highly expressed in ovarian granulosa cells (GCs) from young mice and its expression was significantly decreased in aged GCs. siRNA mediated inhibition of *HAT1* in GCs decreased the polar body extrusion rate, and increased meiotic defects and aneuploidy in oocytes (Chen et al, 2023). Together, these results illustrate the essential role of *Sj*HAT1 in regulating reproductive organ development and egg-laying processes in schistosomes.

Although our *in vitro* results demonstrated significant anti-schistosomal activity via *SjHAT1* knockdown, it is generally acknowledged that this observed efficacy *in vitro* does not always translate in corresponding parasiticidal effects *in vivo* (Patra et al, 2013). In our case, however, *SjHAT1* RNAi *in vivo* caused a 57.5% reduction (*p*<0.0001) in worm burden (Fig. 5A) and 65.9% reduction (*p*<0.0001) in pairing rate (Fig. 5B). Furthermore, the intervention markedly inhibited the developmental progression and maturation of female worms (Figure 5C), with 32.9% arrested at the juvenile stage. This developmental impairment consequently led to compromised oviposition capacity in females. The liver egg deposition was reduced by 79.2% after interference of *SjHAT1*, which in turn significantly attenuated the pathological damage to the mice (Fig. 5F-H). It is well-known that schistosomes are dioecious, with separate male and female sexes (Cort, 1921). The sexual development of female schistosomes is entirely dependent on pairing with a male schistosome (Severinghaus,1928). In the absence of males, the ovaries and vitellaria of females remain in a primordial state, rendering immature females incapable of laying eggs (Clough, 1981). Further, it has been shown that physical contact with a male worm, and not insemination, is sufficient to induce female development and the production of viable eggs (Wang et al, 2019). Chen et al (2022) identified that β-alanyl-tryptamine (BATT), a non-ribosomal peptide produced by male schistosomes is responsible for inducing female sexual development and egg reproduction. Crucially, exogenous BATT alone is sufficient to induce egg laying in immature female parasites (Chen et al, 2022). Thus, the reduced male-female interaction, primarily resulting from the decreased male worm population following *SjHAT1* knockdown, leads to delayed development of female worms. Additionally, functionally impaired male worms exhibit deficient BATT synthesis, which is essential for triggering maturation and oviposition in females. Moreover, similar phenotypic impairments, including testicular hypoplasia, ovarian degeneration and vitellaria atrophy, were observed in reproductive organs of worms recovered from *SjHAT1* RNAi-treated mice *in vivo* (Fig. 5D). These *in vivo* results demonstrate that *Sj*HAT1 critically regulates the growth, pairing and oviposition in *S. japonicum*, suggesting that *Sj*HAT1 is a novel druggable target.

Another striking result from our RNA-seq data is the significant downregulation of *multidrug resistance-associated protein 4* (*MRP4*, KSF78_0000799) in females and *aromatic L-amino acid decarboxylase* (*AADC*, KSF78_0006508) in males. AADCs are the enzymes catalyzing the decarboxylation of aromatic amino acids (such as tryptophan and tyrosine) to generate monoamines (such as tryptamine and dopamine), which play diverse physiological and biosynthetic roles in living organisms (Han & Shin, 2022). It has been reported that in planarians, AADC collaborates with nonribosomal peptide synthetase (NRPS) to catalyze the synthesis of BATT (Issigonis, et al, 2024). BATT is an essential pheromone that stimulates the maturation of the female reproductive system and promotes egg-laying. Furthermore, *in vitro* experiments demonstrate that exogenous BATT can replace male worms to induce female reproductive system development and egg production (Chen et al, 2022). Knockdown of *AADC* in sexual planarians resulted in ovary ablation and loss of accessory reproductive organs, including vitellaria, oviducts, sperm ducts, and gonopore (Khan & Newmark, 2022). Previous studies have found that *AADC* is specifically highly expressed in male *S. japonicum* and is predominantly localized on the inner surface of the gynecophoric canal (Wang et al, 2017), which exhibits spatial overlap with the distribution of *SjHAT1* in male worms. Therefore, we have reason to infer that targeted downregulation of *SjHAT1* suppresses the expression of *AADC*, thereby blocking the synthesis and secretion of the key pheromone BATT in male worms, thus inhibiting the development of the female reproductive system and oviposition. However, how is the BATT signal transduced to female worms? Existing studies have shown that signaling molecules such as linoleic acid, epidermal growth factor, and insulin-like growth factor can enter schistosomes through their excretory system, activating internal signaling pathways (Kusel et al, 2009). ATP binding cassette (ABC) transporters are the main components of the excretory system. Multidrug resistance-associated proteins (MRPs) are members of ABC transporters. Knockdown of *Sm*MRP1 in *S. mansoni* adults disrupts egg production and mitigates liver pathology (Kasinathan et al, 2011). Our RNA-seq data also revealed a significant downregulation of *multidrug resistance-associated protein 4* (*MRP4*) in female worms, indicating that BATT pheromone might be transduced to females through the MRPs. Taken together, we hypothesize that downregulation of *SjHAT1* inhibits the expression of key enzymes such as *AADC* in male worms, blocking the synthesis and excretion of BATT signals in males; by suppressing the ABC transporter system in female worms, it further hinders BATT signal transduction, thereby suppressing the development of the female reproductive system and oviposition. Verification of this hypothesis could provide new strategies for the prevention and control of schistosomiasis, as well as novel targets for the development of anti-schistosomal drugs.

In conclusion, we have characterized *Sj*HAT1 as a novel anti-schistosomal drug target. It is predominantly expressed in the vitellaria of female worms and in the parenchyma adjacent to the gynecophoral canal of male worms. Knockdown of *SjHAT1* impairs worm viability, development and reproduction both *in vitro* and *in vivo*. Furthermore, we hypothesize that dual-sex regulatory mechanisms may play a key role in the process of schistosome female egg-laying. Developing specific *Sj*HAT1 inhibitors could yield novel compounds to control schistosomiasis. Future studies on the downstream mechanisms of *Sj*HAT1 will improve our understanding of sexual maturation and oviposition of schistosomes.

## 4. Experimental Section/Methods

### Ethics statement

All animal experiments were conducted in strict accordance with the recommendations in the Guide for the Care and Use of Laboratory Animals of the National Institutes of Health. All efforts were made to relieve the suffering of the experimental animals. The protocol (including mortality aspects) was approved by the Institutional Animal Care and Use Committee (IACUC) of Soochow University (Permit Number: 2007-13).

### Animals and parasites

Female C57BL/6 mice (6-8 weeks old, 20.0 ± 5 g) were provided by the Center for Experimental Animals at Soochow University (Suzhou, China). All mice were raised under specific pathogen-free (SPF) conditions with a controlled temperature of 22 C°, humidity of 55%, and photoperiod (12 h light, 12 h dark). Each mouse was transcutaneously infected with 60 ± 2 *Schistosoma japonicum* cercariae (Chinese mainland strain) shedding from snails (*Oncomelania hupensis*), which were provided by the Institute of Schistosomiasis Control in Jiangsu Province (Wuxi, China). *S. japonicum* worms were recovered from the infected mice by perfusion through the hepatic portal vein (Duvall & DeWitt,1967). Three worm pairs were cultured at 37_°_C in 5% CO_2_ in 1 ml m-AB169 (1640) medium in a 24-well plate. M-AB169 (1640) is a novel medium that maintains the mature reproductive organs and continuous oviposition of adult female *S. japonicum* (You et al, 2024). The formulation of m-AB169 (1640) consists of RPMI-1640 (Thermo Fisher Scientific, USA), 5% V/V Schneider’s Insect Medium (Merck Millipore, Germany), 10% V/V Fetal Bovine Serum (Solarbio Life Science, Shanghai, China), 1 × antibiotic/antimycotic (100 ×, Solarbio Life Science, Shanghai, China) 0.5 × MEM Vitamin Solution (100 ×, Thermo Fisher Scientific, USA), 0.2 μM triiodothyronine (MCE, USA), 1 μM hydrocortisone (MCE, USA), 1 μM serotonin (MCE, USA), 0.5 μM hypoxanthine (MCE, USA), 8 μg/mL bovine insulin (APExBIO, USA), 0.5 g/L lactalbumin hydrolysate (Sigma-Aldrich, USA), 0.5 g/L casein hydrolysate (Merck Millipore, Germany), 200 μM ascorbic acid (Merck Millipore, Germany), 10 mM HEPES (Thermo Fisher Scientific, USA) and 0.2% V/V bovine red blood cells (Sbjbio Life Science, Nanjing, China). The medium was changed every other day.

### Rapid amplification of cDNA ends (RACE)

Based on the predicted sequences (GenBank: FN318824.1) of *SjHAT1* in the NCBI database, an intermediate fragment was amplified by PCR. Subsequently, gene-specific primers (GSP) and nested gene-specific primers (NGSP) were designed from the intermediate fragment by Primer Premier 6 (Table EV3). RACE was performed with SMARTer RACE cDNA Amplification Kit (Clontech, USA) to obtain the terminal sequences. The purified amplification products were directionally cloned into the pRACE vector using the In-Fusion Cloning System. The resultant recombinant plasmids were subjected to bidirectional Sanger sequencing with M13 universal primers, and the obtained sequences were computationally assembled to determine the full-length cDNA of *SjHAT1*. The open reading frame (ORF) region was analyzed using the ORF Finder tool in the NCBI database, yielding a single complete ORF and its corresponding amino acid sequence.

### Multiple sequence alignment and phylogenetic analysis

Homologous proteins of HAT1 across species were retrieved from the UniProtKB/Swiss-Prot database. Multiple sequence alignment was performed with MAFFT v7.520. Putative conserved protein motifs were identified via MEME Suite. The phylogenetic tree construction via the Maximum Likelihood method in MEGA11 with 1000 bootstrap replicates for node support evaluation, and subsequently visualized using Chiplot (https://www.chiplot.online).

### Quantitative real-time PCR (qRT-PCR)

TRIzol Reagent (Invitrogen, USA) was used to extract total RNA from different developmental stages of *S. japonicum*, including eggs, cercariae, 14-day schistosomula, 21-day, 28-day, and 35-day adult male and female worms. Subsequently, 1 μg total RNA was used for first-strand cDNA synthesis using the RevertAid First-Strand cDNA Synthesis Kit (Thermo Fisher Scientific, USA). The cDNA was used as a template for qRT-PCR with ChamQ Universal SYBR qPCR Master Mix (Vazyme, China) and 0.4 µM forward and reverse primers. Amplification was performed on CFX96 Touch Quantitative Real-Time PCR System (Bio-Rad, USA) under standardized conditions: 95℃ for 3 min, followed by 40 cycles of 95℃ for 10 s, 60°C for 30 s, 72℃ for 10 s; melt curve analysis: 65℃ for 5 s, 95℃ for 5 s. The qPCR primers (Table EV3) were designed with Primer Premier 6.0 software and validated through melting curve analysis. Relative quantification was performed using the 2^-Δ^ ^ΔCt^ method with *S. japonicum* 26S proteasome non-ATPase regulatory subunit 4 (*SjPSMD4*, GenBank No. FN320595.1) as the internal reference.

### Fluorescence *in situ* hybridization (FISH)

Males and females recovered from the infected mice at 35 days postinfection (dpi) were fixed in 4% paraformaldehyde at room temperature for 4 h, washed with PBSTX (1×PBS, 0.3% Triton X-100), and treated with a permeabilization solution (50 mM DTT, 0.1% SDS, 1% NP-40 in 1×PBS) at 37℃ for 10 min. Samples were dehydrated in 100% PBSTX and graded methanol (25%, 50%, 75%, 100%), bleached in 6% bleach solution (30% H_2_O_2_ diluted in 100% methanol) under direct light at 4℃ for 20 h, and rehydrated in reverse methanol gradients. Rehydrated samples were treated with proteinase K solution (1 μg/ml proteinase K, 0.1% SDS in 1×PBS) at 37℃ for 20 min, re-fixed in 4% paraformaldehyde for 10 min, and incubated with autofluorescence quenchers (Servicebio, China) for 30 min. A CY3-labeled oligonucleotide probe (Table EV3) synthesized by Huzhou Hippo Biotechnology Co., Ltd was hybridized at 56℃ for 4 h. Nuclei were counterstained with DAPI, and samples were mounted with anti-fade medium. Visualization was performed on a Leica TCS SP8 Laser Scanning Confocal Microscope (Leica, Germany) using 10× objective.

### Synthesis of dsRNA

The dsRNA templates for *SjHAT1*-dsRNA and *GFP*-dsRNA were amplified from the pGEX-4T-1-*Sj*HAT1 maintained in our lab and commercial pET-28a-GFP synthesized by Sangon Biotech (Shanghai, China), respectively, using T7 promoter-containing primers (Table EV3). The size of amplification products was verified by 1% agarose gel electrophoresis, and the identity was confirmed through DNA sequencing. Purified PCR products were employed to synthesis single-stranded RNA using the MEGAshortscript™ T7 Kit (Invitrogen, USA), following the manufacturer’s protocols. To synthesize dsRNA, a transcription reaction mixture containing equimolar sense and antisense ssRNA strands was denatured initially in a 75 ℃ water bath for 5 min, followed by annealing at room temperature for 30 min. The resulting dsRNA was subsequently treated with TURBO DNase (Invitrogen, USA) and RNase If (New England Biolabs, USA) to remove residual DNA templates and single-stranded RNA contaminants. After phenol/chloroform extraction and ethanol precipitation, the dsRNA was resuspended in DEPC-treated water. The concentration of dsRNA was quantified using a NanoDrop 2000c Spectrophotometer (Thermo Fisher Scientific, USA) and adjusted to 3.0 μg/μL. The integrity and size of dsRNA were verified by electrophoresis on 1% agarose gels, and the products were stored at −80 ℃.

### SjHAT1-RNAi in vitro

*S. japonicum* paired adults (35 dpi) were obtained and cultured in m-AB169 (1640). *Green fluorescent protein* (*GFP*) dsRNA served as the negative control for all RNAi experiments. D1 represents the first day of the experiment. Paired adults were treated with 30 μg/mL dsRNA of *GFP* or *SjHAT1* on D1, D3 and D5 with fresh medium. After daily viability assessments from D1 to D7, worms were collected for stereomicroscopy analysis (Nikon, JPN), and the viability score was assigned as described previously (Santiago Ede et al, 2014), based on the changes in mobility and general appearance. Briefly, each worm was assigned a viability score ranging from 0 to 3. Score 3: worms exhibited the highest activity level as observed in the control group, moving actively and smoothly with transparent bodies; Score 2: worms showed whole-body movement but appeared stiff and slow, with translucent bodies; Score 1: partial movement was observed with opaque body appearance; Score 0: worms remained contracted without resuming movement, considered dead. Concurrently, eggs in the culture medium were collected and enumerated on D3, D5, and D7.

### SjHAT1-RNAi in vivo

Nine *S. japonicum* infected mice were randomly divided into three groups, *SjHAT1*-dsRNA group, *GFP*-dsRNA control group, and blank control group, with three mice per group. Ten micrograms of *SjHAT1* or *GFP* dsRNA were injected into the tail vein for interference at 14 dpi, 18 dpi, 22 dpi, 26 dpi, and 30 dpi. An equal volume of 0.7% saline was injected into each mouse in the blank control group at the same time point. Following euthanasia at 35 dpi, adult worms were collected for both RNAi efficacy assessment and phenotypic analysis, while liver tissues were analyzed for egg quantification (see the section “Liver egg counting”) and histopathology (see the section “Histopathological assessment”). The experiments were repeated thrice.

### Efficacy of *SjHAT1*-RNAi

After treatment with *SjHAT1* dsRNA, transcriptional levels of *SjHAT1* of adult worms were quantified by qRT-PCR performed as previously described. Protein expression levels of *Sj*HAT1 were assessed by Western blotting. To prepare the adult worm soluble protein extracts, phosphate buffer saline precooled on ice containing 20 adult worms was homogenized by sonication (30% power, 5s on, 5s off, 2 min, Q700 Sonicator, USA), and centrifuged for 10min at 13,000 g, 4℃. Protein concentrations were determined by BCA assay. Worm extracts (20 μg protein) were separated on 10% SDS-PAGE gels and transferred to 0.45 μm PVDF membranes (Millipore, Germany). The membranes were blocked with rapid blocking buffer for 15 min at room temperature, then washed with TBST and incubated overnight with rabbit anti-*Sj*HAT1 polyclonal antibody used at 1:500 dilution in TBST at 4℃. After washing with TBST, membranes were incubated for 1 h at room temperature in TBST containing HRP-conjugated goat anti-rabbit IgG (CST, USA) at a dilution of 1:2000. After washing in TBST, the membrane was developed with BeyoECL plus solution (Beyotime Biotech, Shanghai, China) and imaged using Tanon 5200 (Tanon, Shanghai, China).

### Morphological observations of schistosomes after interference of *SjHAT1*

Adult male and female worms were fixed in 4% paraformaldehyde at room temperature overnight, gently compressed between glass slides for 1 h. Specimens were stained with hydrochloric carmine for 2 h, destained in 70% ethanol containing 1% HCl for 20 min, and dehydrated through an ethanol series (80%, 90%, 95%, 100%). After equilibration in ethanol-xylene (1:1) and clearing in 100% xylene, worms were mounted with neutral balsam on glass slides for morphometric and morphological analyses. The morphologies of reproductive organs and germ cells of *S. japonicum* were observed using a Leica TCS SP8 laser scanning confocal microscope (Leica, Germany) with a 63×oil immersion objective.

### Scanning electron microscopy (SEM)

For SEM analysis, worms were fixed in 1% osmium tetroxide and dehydrated in graded ethanol. After that, the specimens were dried for approximately 30 min. Then the worm samples were mounted on aluminum stubs, coated with gold, and examined with a Hitachi-S4700 scanning electron microscope (Chiyodaku, Japan).

### Liver egg counting

0.5g of mouse liver was weighed and incubated overnight in 5% sodium hydroxide (NaOH) at 37°C and 220 rpm to obtain a homogeneous solution. Ten μL of the above solution was observed and counted under an ordinary upright microscope. Each sample was counted 10 times, and the average was calculated. Each group had at least three replicates. The number of liver eggs (EPG) was calculated using the formula: EPG = average×1000/ 0.5

### Histopathological assessment

The liver of each mouse was washed with PBS (pH 7.4) and fixed with paraformaldehyde. After dehydration, the tissues were embedded in paraffin. Sections 4-μm-thick were stained with hematoxylin and eosin for granuloma analysis. The areas of granulomas surrounding single egg were observed at 200× magnification.

### Statistics

Data were analyzed with GraphPad Prism or Excel, expressed as means ± SD, and tested for statistical significance using either ANOVA or *t*-tests, as noted in the figure legends.

### RNA sequencing

Worms incubated with *SjHAT1*- and *GFP*-dsRNA for 7 days were collected and washed three times with 1×PBS (pH 7.4). After quick freezing in liquid nitrogen, the samples were sent to SeqHealth Technology Corporation (Wuhan, China) for RNA extraction, library construction and sequencing. RNA integrity was validated using the LabChip GX Touch system (PerkinElmer, USA), followed by fluorometric quantification via Qubit 3.0 Fluorometer (Thermo Fisher Scientific, USA). Libraries were constructed from 500 ng total RNA using the KC mRNA Library Prep Kit (Seqhealth, China) according to the manufacturer’s protocol and sequencing was performed on the DNBSEQ-T7 platform (MGI, China).

### Analysis of gene expression profiles and differential expression genes

The FASTQC program (https://www.bioinformatics.babraham.ac.uk/projects/fastqc/) performed quality control on Raw data. FASTP v0.23.2 was used to remove low-quality reads and trim adapter sequences to generate clean reads. Clean reads were aligned to the *S. japonicum* genome (ASM2146165v1) using STAR v2.7.6a, followed by exonic read quantification with featureCounts (Subread-1.5.1) and RPKM normalization for gene expression profiling. Differential expression analysis was performed using the edgeR package (v3.40.2) with thresholds of fold change (FC)>1.5 or < 1.5 and *p*<0.05. Statistically significant GO terms and KEGG pathways enriched in differentially expressed genes (DEGs) were identified using KOBAS v2.1.1 with a significance threshold of *p* < 0.05. Transcriptional validation of target genes was performed through qRT-PCR as previously described, with primer sequences listed in Table EV3.

## Data Availability Statement

All data are available in the main text or the supplementary materials. The accession number of full-length *SjHAT1* cDNA in NCBI database is PQ778043. The raw RNA sequencing data of male and female *S. japonicum* worms treated with *GFP*-dsRNA or *SjHAT1*-dsRNA have been deposited in the Sequence Read Archive (SRA) database under the accession number PRJNA1313613.

## Acknowledgements

We thank Professor Xing-Quan Zhu for polishing and revising the manuscript. The infected *Oncomelania hupensis* snails were provided by the Institute of Schistosomiasis Control in Jiangsu Province (Wuxi, China). This work was supported by National Natural Science Foundation of China (No. 81902083, No. 82172294), Priority Academic Program Development of Jiangsu Higher Education Institutions (No. YX13400214), and Key Laboratory of Basic Research on Regional Diseases (Guangxi Medical University), Education Department of Guangxi Zhuang Autonomous Region (No. GXQYJB2024001)

## Author Contributions

JX and YXW contributed equally to this work. JX and CMX conceived and designed the study. YXW and JX performed the experiments. JX prepared the manuscript. PH, YNZ and HS contributed to data analyses.

## The Paper Explained Problem

Schistosomiasis, a neglected tropical disease caused by *Schistosoma* species, remains a major public health challenge in low-income regions. In the absence of an effective vaccine, praziquantel (PZQ) remains the primary therapeutic agent for the treatment and control of schistosomiasis in most developing countries. However, repeated and prolonged mass drug administration of PZQ has raised growing concerns about the emergence of drug resistance. Consequently, developing alternative theraputic approches and identifying novel drug targets have become critical priorities in addressing this debilitating parasitic disease.

## Result

Our work focused on discovering novel anti-schistosomal drugs, and we have developed a novel praziquantel derivative, DW-3-15 (patent no. ZL201110142538.2), which demonstrated consistently potent broad-spectrum schistosomicidal activity *in vitro* and *in vivo*, with especially high efficacy against juveniles. Further studies revealed that its therapeutic effect is mediated through the downregulation of *Schistosoma japonicum* histone acetyltransferase 1 (*Sj*HAT1). Here, we clone the full-length *SjHAT1* cDNA (NCBI accession no. PQ778043). The encoded protein retains all catalytic residues critical for acetyltransferase activity despite sharing only ∼34% sequence identity with mammalian HAT1 orthologs. Phylogenetic analysis place *Sj*HAT1 in a distinct evolutionary clade. Expression profiling across developmental stages reveals significant stage- and sex-specific differences. The location of *SjHAT1* is predominantly in the vitellaria of female worms and in the parenchyma near the gynecophoral canal of male worms. *In vitro* RNA interference of *SjHAT1* significantly reduces the adult worm viability and female oviposition. *In vivo*, tail-vein injection of *SjHAT1*-dsRNA in infected mice leads to a 57.5% reduction in adult worm burden, a 65.9% reduction in worm pairing and a 79.2% reduction in liver tissue egg burden. RNA-sequencing analysis reveals that *SjHAT1* knockdown disrupted β-alanyl-tryptamine (BATT) pheromone signaling via downregulation of aromatic *L-amino acid decarboxylase* (*AADC*) in males and *multidrug resistance-associated protein 4* (*MRP4*) in females.

### Impact

In this study, we identify that *Sj*HAT1 as a promising anti-schistosomal drug target. Moreover, our results demonstrate that *Sj*HAT1 plays a critical role in regulating the synthesis of BATT, an essential male pheromone required to stimulate female reproductive development. Knockdown of *SjHAT1* downregulates *AADC*, thereby blocking the transmission of BATT signal to female worms. Concurrently, downregulation of female-specific *MRP4* impairs BATT signal transduction in female worms, further suppressing oviposition. The dual-sex regulatory mechanisms identified here will provide new insights into the sexual biology of schistosomes and could inspire novel strategies for schistosomiasis control.

## Expanded View Figure legends

**Figure EV1.**
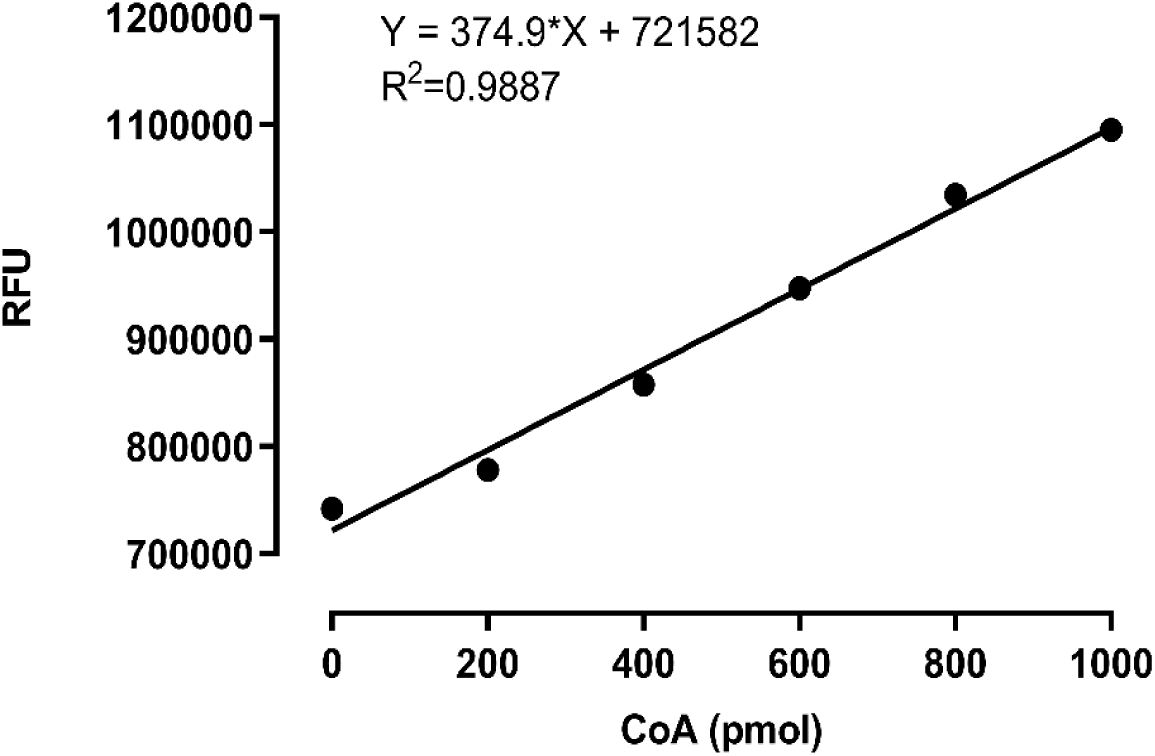
CoA Standard curve. The standard curve for CoA content was established using the reference standard from the Histone Acetyltransferase Activity Assay Kit (Fluorometric, Catalog No. AB204709, Abcam, USA), The regression equation is Y = 374.9X + 721,582, R^2^=0.9887, where the vertical axis represents the relative fluorescence units (RFU) and the horizontal axis corresponds to the CoA content (pmol).

**Figure EV2.**
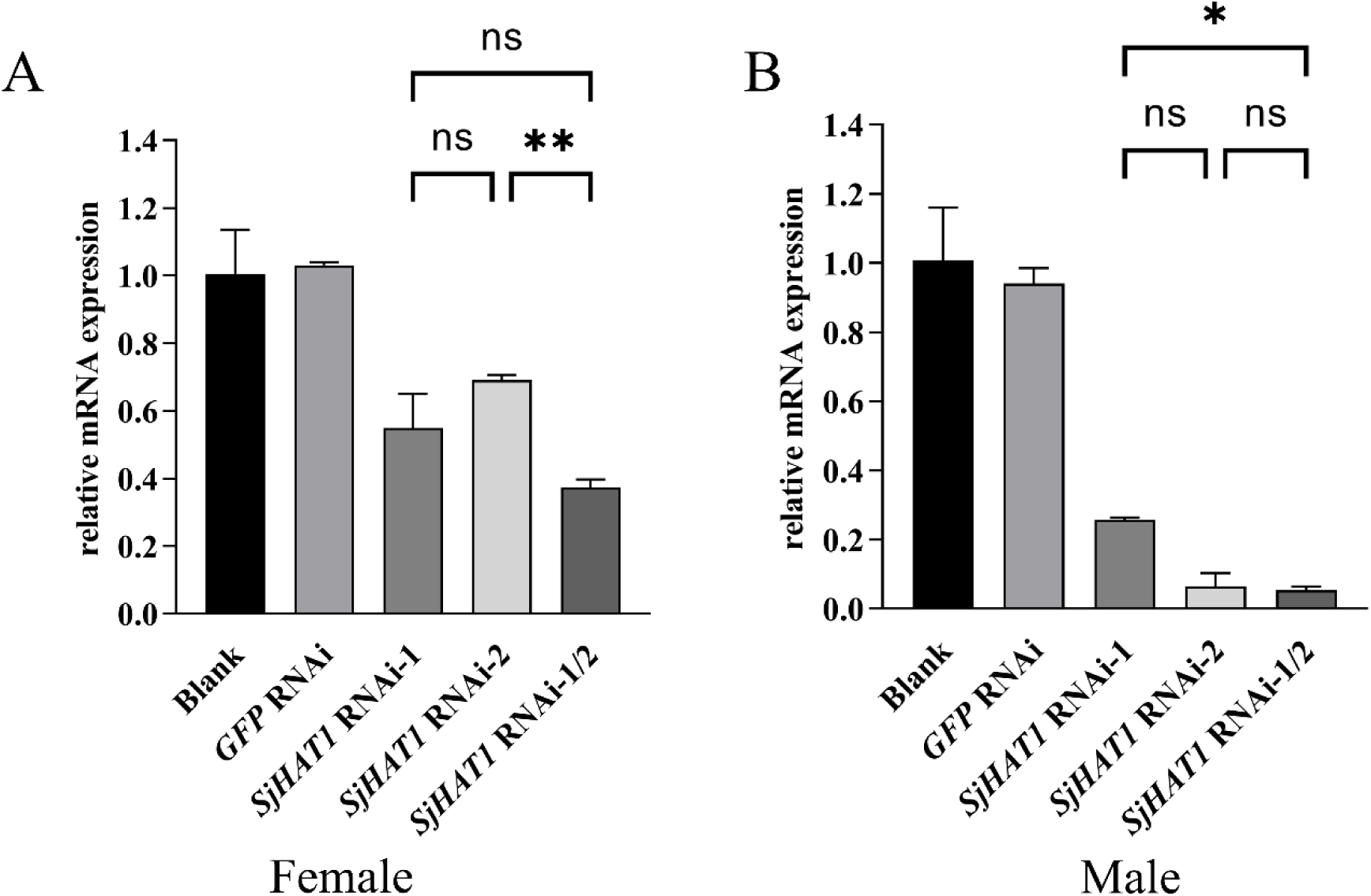
Efficiency of RNA interference for *SjHAT1*-dsRNA. Adult female and male worms with good activity were added to the 24-well plate with five worm per well, different kinds of *SjHAT1*-dsRNA were added for interference on the first, third, and fifth days. Each group had three replicate wells for each biological experiment, with three biological replicates. DEPC-treated water served as the blank control, and *green fluorescent protein* (*GFP*) dsRNA was used as the negative control. (**A**) Comparison of RNA interference efficiency among *SjHAT1*-dsRNA1 (30 μg/mL), *SjHAT1*-dsRNA2 (30 μg/mL), and *SjHAT1*-dsRNA1/2 (15 μg/mL *SjHAT1*-dsRNA1+15 μg/mL *SjHAT1*-dsRNA2) in female worms. (**B**) Comparison of RNA interference efficiency among *SjHAT1*-dsRNA1, *SjHAT1*-dsRNA2, and *SjHAT1*-dsRNA1/2 in male worms. Error bars indicate standard deviation (SD). Statistical significance is denoted as follows: ‘ns’, not significant, \**p* < 0.05, \*\**p* <0.01 (One-Way ANOVA).

**Figure EV3.**
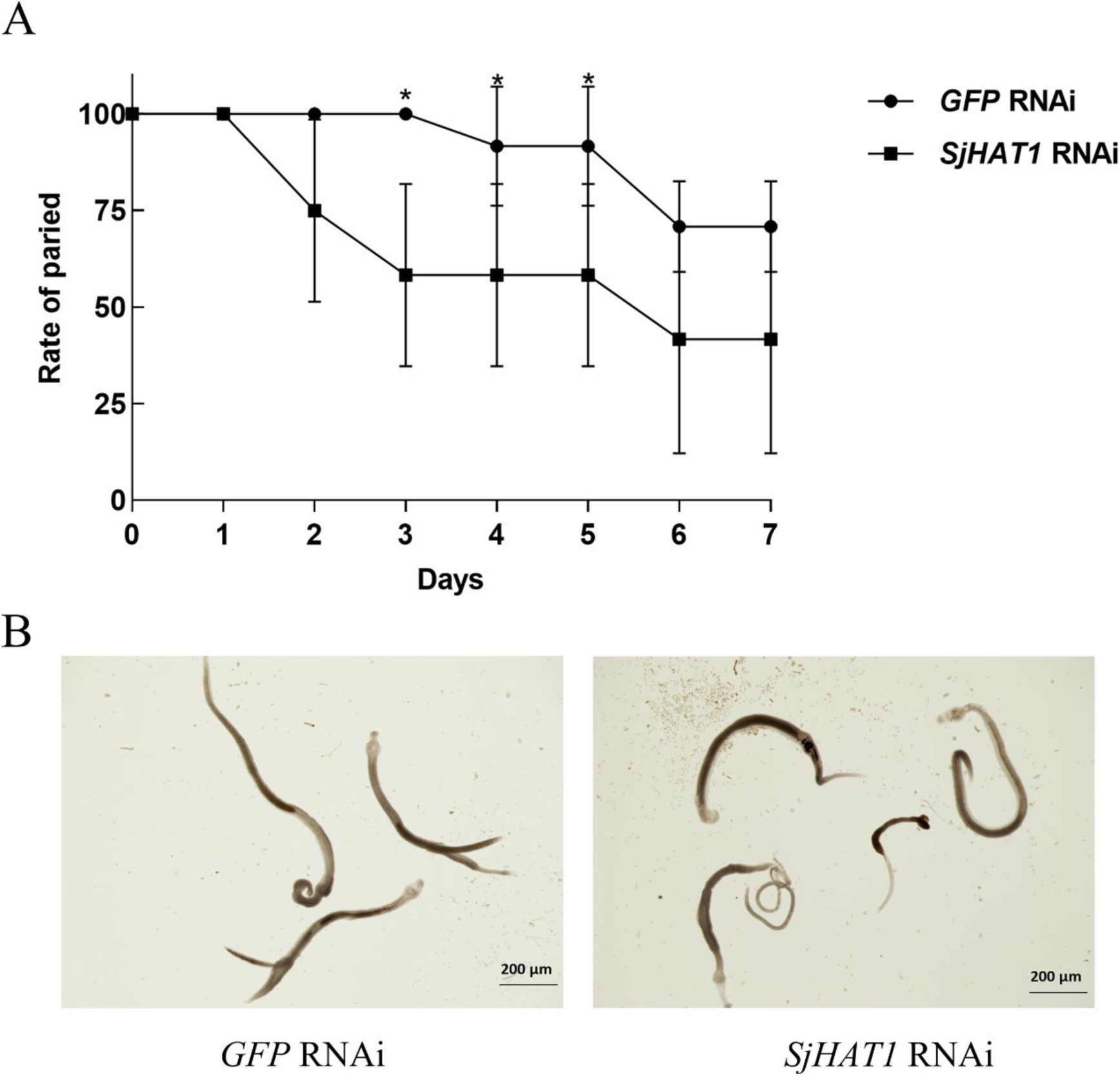
Effect of *SjHAT1*-dsRNA on worm pairing. Adults with good pairing activity were incubated in the 24-well plate with three pairs per well. Each group had two replicate wells for each biological experiment, with four biological replicates. **(A)** Effect of *SjHAT1*-dsRNA on pairing rate. **(B)** Observation under the light microscope on D7 after treatment with *GFP*-dsRNA and *SjHAT1*-dsRNA. Error bars indicate standard deviation (SD). \**p* < 0.01 (Two-Way ANOVA).

**Figure EV4.**
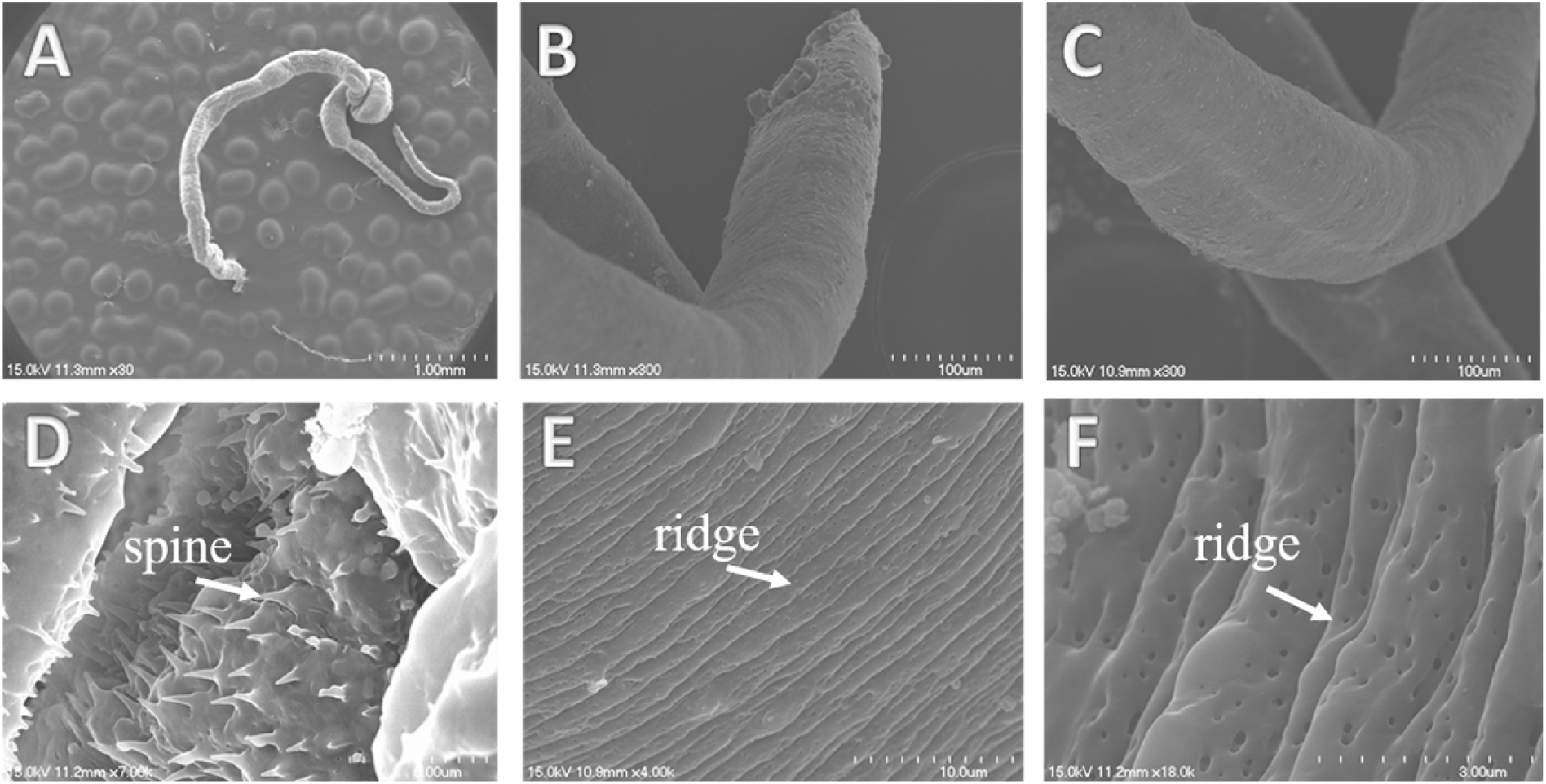
Scanning electron micrographs of the tegument of *S. japonicum* females in the control group after 3 days incubation. (A) Gentle panorama of a female worm. (B) Anterior part of the female worm. (C) Mid-portion of the female worm. (D) Spines in the ventral sucker of the female. (E)-(F) Transverse ridges on the tegument of the female worm.

**Figure EV5.**
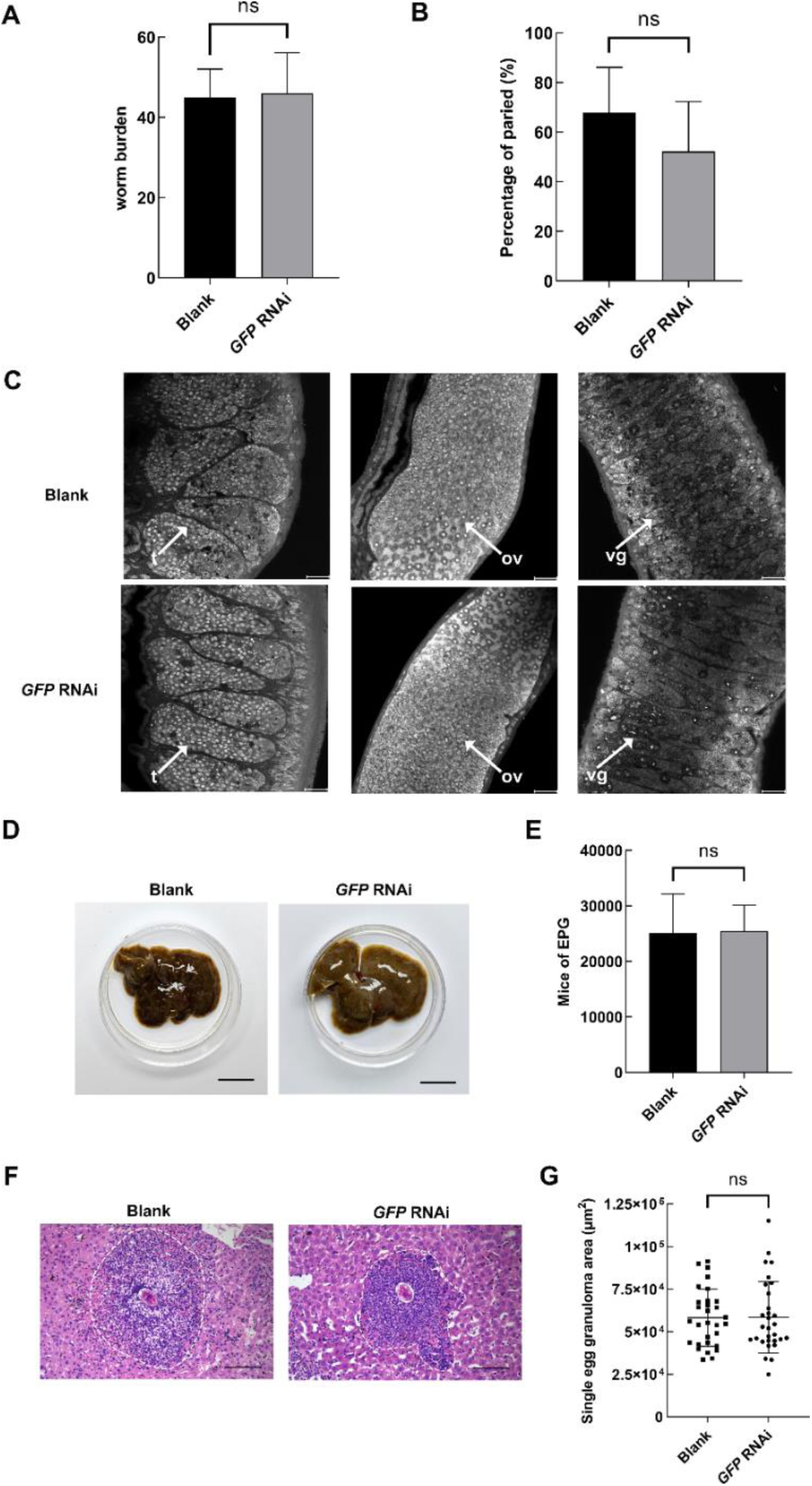
*GFP* RNAi has no effect on worm survival, worm pairing and oviposition *in vivo*. On the 14th, 18th, 22th, 26th and 30th days postinfection, *GFP*-dsRNA was injected through the tail vein, and the hepatic portal vein was perfused on the 35^th^ day. (A) Worm burden of the parasites recovered at 35 dpi in Blank and *GFP* RNAi groups. (B) Comparation of the pairing rates of recovered worms between blank control group and *GFP* RNAi group. (C) Observation of the reproductive organs of worms after *in vivo* treatment. t, testis; ov, ovary; vg, vitelline gland. Scale bars: 20 μm. (D) Gross observations of the mouse liver from the blank and *GFP* RNAi group. Scale bars: 1 cm. (E) Egg count per gram of liver comparation between the blank and *GFP* RNAi groups. (F) Histological assessment of mouse liver by H&E staining. Scale bars: 100 μm. (G) Statistical analysis of the size of egg granuloma area after *in vivo* treatment, with 30 egg granulomas per group. Standard deviation (SD) is shown in the error bars. “ns”, not significant (Student’s *t*-test).

**Table EV1.** Quantification of r*Sj*HAT1 enzymatic activity.

| group | sample quality<br>(ng) | RFUs | CoA production<br>(pmol) | Acetyltransferase<br>(HAT) activity<br>($\mu\text{U}/\mu\text{g}$) |
| --- | --- | --- | --- | --- |
| Blank control | 0 | 618926±25398.2 | 0 | 0 |
| Negative<br>control | 50 | 625270.5±1888.5 | 0 | 0 |
| rSjHAT1 | 50 | 1224562.5±5854.6 | 1341.6±15.6 | 447.2±5.2**** |

\*\*\*\* $p < 0.0001$ (One-Way ANOVA).
HAT activity ( $\mu\text{U}/\mu\text{g}$ ) = Amount of CoA from standard curve (pmol) / [reaction time (min) × original sample quality added into the reaction well ( $\mu\text{g}$ )]. In this experiment, the reaction time was set to 60 min.

**Table EV2.** The number of mature and immature female worms recovered on 35^th^ day post RNA interference treatment with *GFP*-dsRNA and *SjHAT1*-dsRNA.

| group | Number of mature female worm | Number of immature female worm |
| --- | --- | --- |
| *GFP* RNAi | 138 | 32 |
| *SjHAT1* RNAi | 47 | 23 |

Chi-square test: $\chi^2(1) = 5.528, p = 0.0187$

**Table EV3.**
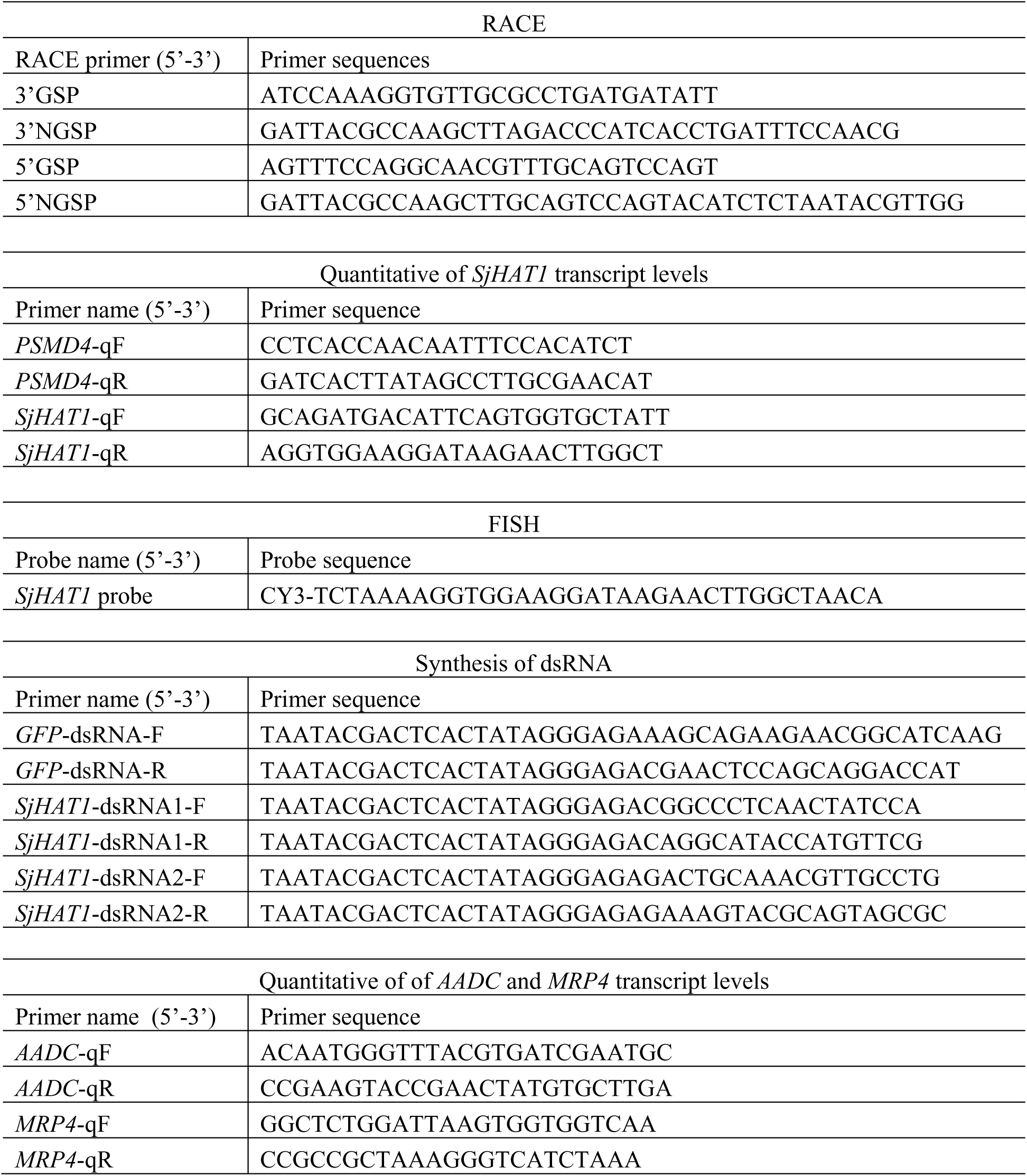
Primers list.

